# Rapid Rescue: Spatial Storage Effect Facilitates Evolutionary Rescue in Rapidly Changing Environments

**DOI:** 10.1101/2025.02.06.636948

**Authors:** Eve Nancy Rowland, Davorka Gulisija

## Abstract

The storage effect is a plausible natural mechanism that generates balanced genetic polymorphism in varying environments. Balanced polymorphism may facilitate evolutionary rescue, promoting the persistence of populations otherwise destined for extinction. However, it is unknown whether the storage effect can be established in small populations whose size is allowed to vary, and if so, whether it will lead to evolutionary rescue. In this study, we investigate if the spatial storage effect emerges and facilitates evolutionary rescue across small populations of variable sizes that inhabit heterogeneous temporally varying environments and exchange migrants. We use an eco-evolutionary model to examine the phenomenon under a wide set of conditions, including the magnitudes and periods of temporal variation, habitat harshness, migration rates, the degrees of spatial heterogeneity, and increasing fitness oscillations over time, all within the framework of the logistic population growth model. We find that the storage effect emerges and that it increases the persistence of populations in harsh temporally varying habitats beyond levels driven by migration alone under source-sink dynamics. This mechanism, which we call “rapid rescue” broadens the known conditions for population persistence in the face of sudden environmental change.

## Introduction

Many populations experience temporally varying selection resulting from seasonal, periodic, or random environmental fluctuations [1–4]. Some populations experience changes in the patterns of temporal environmental variation due to global climate change [5,6], anthropogenetic habitat alteration [7–9], or invasions into novel habitats [10]. Global climate change, in particular, is expected to increase the magnitude of environmental variation via extreme temperature [11] and precipitation events [12,13]. These changes may become a major driving force of evolution on contemporary timescales [14]. In response to new varying regimes, populations can become extinct, expand their range, or adapt to rapidly changing conditions [15–20]. The latter represents a particular mode of evolution, referred to as rapid evolution [21–24], where populations undergo rapid shifts in allele frequencies on contemporary timescales [25,26]. The conditions that drive rapid evolution in temporally varying environments differ notably from those under traditionally considered gradual evolution [22].

Rapid evolution occurs on ecological time scales [27] and requires notable genetic variation [24,28]. First, rapid evolution occurring on ecological time scales implies that ecological and evolutionary processes jointly impact how populations change over time [29]. Interactions between ecological and evolutionary processes are known to produce specific patterns of genetic change in populations [30–32]. Nonetheless, how such interactions contribute to rapid evolution in temporally varying environments is underexplored. In particular, the studies that examine eco-evolutionary dynamics tend to focus on directional selection [33] or deterministic dynamics [32]. Second, the process of rapid evolution may be contingent on the levels of genetic variation [24,28] since it is more likely to occur if adaptive alleles are already present in a population [34,35]. Such variation may be supplied by a high rate of beneficial *de novo* mutations [36], which is unlikely in small populations facing the risk of extinction [34,37,38]. Therefore, rapid adaptation in small populations might depend on genetic diversity that has been maintained over time, known as balanced polymorphism [39–44]. However, it is unclear how the eco-evolutionary dynamics affect the maintenance of genetic polymorphism in temporally varying environments.

Classical population genetics theory suggests that the maintenance of genetic variation is restricted to heterozygous advantage across environments [44–47] and is unlikely in finite populations [41]. Recent models incorporating forms of the storage effect, originally explored in the context of species coexistence [48], emerged as plausible mechanisms of balancing selection in temporally varying environments [49–56]. The storage effect arises in temporally varying habitats under strong density regulation (competition), when a portion of a population is (partially) protected from selection [48,56]. For example, storage of genetic polymorphism in populations can occur due to partial protection from selection in different life-stages (e.g., seedbanks, long-lived adults, diapause, etc.) [57,58], under phenotypic plasticity [52], under maternal effects [53], or across habitats with a spatially heterogenous magnitude of variation [51]. The latter is referred to as the spatial storage effect and is a mechanism that easily establishes under a wide range of naturally plausible scenarios, including variation of the levels of selection, the rate of migration across the structured population, and the degree of spatial heterogeneity in selection [51]. Moreover, this mechanism has been explored under eco-evolutionary dynamics and has been found to arise when population carrying capacity fluctuates [32], when modeled under deterministic dynamics. However, none of the studies on the storage effect incorporated stochastic changes in population sizes nor allowed for possible extinction. Thus, it remains unclear if the storage effect generates notable genetic diversity under population eco-evolutionary dynamics and if so, how it impacts population persistence amid stochastic perturbations to population size.

Genetic diversity may prevent population extinction [27,37,59]. That is, threatened populations can be “rescued” from extinction by adaptation: a phenomenon known as “evolutionary rescue” [60–66]. The presence of adaptive alleles (i.e., genetic variation) is a limiting factor for evolutionary rescue [67,68]. Rapidly changing conditions may require rapid evolutionary rescue from balanced polymorphism in small populations, given the limited influx of adaptive mutants from *de novo* mutations. In this scenario, a form of the storage effect, such as the spatial storage effect, could maintain sufficient balanced polymorphism needed for rapid evolution as environments shift. Therefore, it is imperative to investigate whether the spatial storage effect can arise in small populations under eco-evolutionary population dynamics, maintain genetic diversity over time, and lead to evolutionary rescue under continuously changing conditions.

An important consideration when investigating evolutionary rescue under the spatial storage effect is population persistence in the absence of evolution. Under the conditions favoring the spatial storage effect, where populations in distinct temporally varying regimes exchange migrants, migration may support population persistence without evolution by supplying individuals through a source-sink dynamic [69,70]. A source-sink dynamic occurs when a population inhabiting favorable conditions (source) supplies migrants to a population in a poor-quality habitat (sink) [69,71,72] thus maintaining population size [73]. As long as the migration rates remain moderate, migration promotes persistence by maintaining connectivity between spatially heterogeneous populations, thus protecting fragmented populations [74]. The main question that arises is whether the spatial storage effect can increase the rates of population persistence to a higher level than migration alone, thus, representing a case of evolutionary rescue.

This study explores whether the storage effect emerges and leads to rapid evolutionary rescue in small populations whose sizes are allowed to vary. We develop a population model of the spatial storage effect [51], arising across two populations experiencing different magnitudes of varying selection and environmental harshness. These populations exchange migrants and undergo size changes according to the logistic growth model. We use forward-in-time computer simulations over a wide range of parameters, including a range of migration rates, the rate of environmental oscillations, the magnitude of varying selection, and the magnitude of mean environmental harshness, generating scenarios that match contemporary ecological processes, including population extinction. We show that the storage effect establishes and facilitates rapid evolutionary rescue in small populations with stochastic changes in size, generating persistence exceeding that under source-sink dynamics in rapidly changing environments. We call this phenomenon “rapid rescue”.

## Model and Methods

### Eco-evolutionary model

We model eco-evolutionary dynamics at a single locus across two populations that occupy distinct habitats and exchange a limited number of migrants, thus generating conditions that favor the spatial storage effect. The populations originate from a larger stable ancestral population that is under drift (or reduced magnitude of varying selection)-mutation balance for the two alleles, ancestral and derived, *a* and *d*. At the onset of temporally varying selection, the initial allele frequencies of the derived and ancestral alleles within new populations are sampled from the equilibrium distribution of allele frequencies from the ancestral population (*Figure S1*). Then, one population experiences temporally varying selection (population V, with size *N*_V,*t*_ at a time *t*), while the other (population R, with size *N*_R,*t*_ at a time *t*) serves as a refuge that is under drift or a reduced magnitude of temporally varying selection, continuing under the ancestral conditions. To facilitate the presentation of the eco-evolutionary model below, we show the symbols used in the study in *Table 1* where *i* indicates population V or R.

**Table 1.** Mathematical symbols and parameters used in the study.

| Syntax | Description |
| --- | --- |
| $N_{i,t}$ | Size of population $i$ at a time $t$ |
| $K_i$ | Carrying capacity of population $i$ |
| $G_{i,t}$ | Growth in number of individuals in population $i$ at time $t$ |
| $h$ | Heterozygosity across populations under the model relative to neutral expectation (model/drift) |
| $p_{i,t}$ | Frequency of ancestral allele in population $i$ at time $t$ |
| $q_{i,t}$ | Frequency of derived allele in population $i$ at time $t$ |
| $P$ | Number of generations in a cycle of varying selection (phase) |
| $s_{\max}$ | Maximum numerator fitness deviation due to varying selection |
| $M_i$ | Migration rate from population $i$ |
| $\delta_i$ | Intrinsic growth rate in population $i$ |
| $S_i$ | Mean environmental effect in population $i$ |
| $\beta$ | Relative increase in the magnitude of fitness oscillations over time |

We simulate eco-evolutionary dynamics across two populations forward in time [75]. During each discrete generation, the populations are subjected to forces of stochastic mutation under a recurrent mutation rate *μ*, stochastic migration, temporally varying selection, and stochastic reproduction in the framework of the logistic population growth model. Migration is reciprocal such that an equal proportion, given by the migration rate (*m*), of each population is randomly selected to be exchanged with one another. The number of migrants from a population (*M_i_*) is thus *mN_i,t_*.

Temporal environmental variation, implemented as a genotype-based relative fitness deviation at time *t* (*s_t_*), in combination with the overall environmental harshness (*S*_i_ = mean environmental effect in population *i*) and population growth rate (*G_i,t_*), impacts the number of individuals of each genotype in each generation such that:

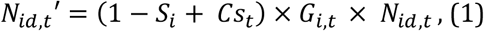

and

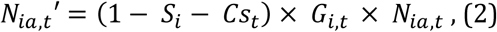

where *N_id,t_* represents the number of individuals carrying the derived allele and *N_ia,t_* represents the number of individuals carrying the ancestral allele in the population *i* at the time *t* prior to density-regulated selection, where *i* = V or R. *S_i_* is a general habitat effect that both alleles experience equally and thus does not induce selective dynamics. *S*_V_ > 0 represents a harsh new habitat for the V population, while the R population resides in a non-harsh habitat with *S*_R_ = 0, consistent with source-sink dynamics [69].

The selection process (non-neutral evolution) is imposed by *Cs_t_* and is such that alleles have opposite effects, so that when an allele is advantageous the other one is deleterious and vice versa, oscillating in effects as the environment changes. Over the cycle of fitness oscillations, both alleles experience the same magnitude and periods of fitness advantages and disadvantages, resulting in alleles being quasi-neutral [76]. In V populations, *C* = 1 always, meaning the population experiences the full magnitude of fitness oscillations. R experiences a lower magnitude of temporal oscillations (0 ≤ *C* < 1) and acts as the refuge needed to establish the spatial storage effect. Therefore, *C* is the portion of relative fitness oscillations in R compared to the V population.

The variation in fitness emulates seasonal oscillations such as those due to changes in temperature, precipitation, or resource availability. The numerator fitness value *s_t_* is generated using the sine function with maximum deviation *s*_max_, such that *s_t_* provides either a selective advantage or disadvantage at time *t* for the derived allele,

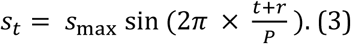

Here *t* represents a generation within an oscillation cycle that is *P* generations long and *r* is a value randomly selected between 0 and *P,* which determines the point within the sine function that the simulation begins. This ensures that the derived allele can randomly start as advantageous or deleterious, depending on the point in the environmental cycle. If *s*_max_ = 0 alleles evolve under neutrality.

The population size changes based on a logistic growth equation with

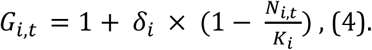

The number of individuals for the two alleles in each population in the next generation are sampled from the Poisson distribution with mean *N*_*ia*,*t*_′ and *N*_*id*,*t*_′ during the stochastic reproduction step.

#### Increasing the magnitude of varying selection over time

To simulate the increasing magnitude of fitness oscillations over time, we modify the numerator fitness deviation such that it is an increasing function of the effect under the constant magnitude of starting s_max_, *s_cm,t_*, generated using equation (3):

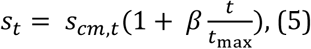

where *β* is the total relative increase in the magnitude of fitness oscillations over time.

#### Measuring genetic diversity and persistence

The expected heterozygosity in a population (*i*) at the time *t* is

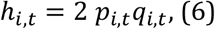

where *p_i,t_* is the frequency of the ancestral allele, *a*, and *q_i,t_* is the frequency of the derived allele, *d*, in a population *i* at time *t*. The weighted heterozygosity at the time *t* across V and R is, thus,

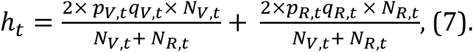

We report expected heterozygosity relative to drift over time and populations,

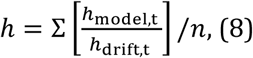

where n is the number of replicate simulations (populations). We define the relative heterozygosity *h >* 1.05, as an indicator that the storage effect has been established. As both heterozygosity in residents and immigrants play a role in rapid adaptation in V, we measure heterozygosity at the level of the meta-system. The weighting of local levels of heterozygosity in (7) assures that we estimate a probability of choosing two different alleles within a population and not across populations, which signifies differentiation (see Gulisija& Kim, 2015 [51]).

#### Measuring Extinction

Each simulation run continues until the maximum allotted time (100*K_i_* generations) or until V goes extinct. Extinction is defined by the subpopulation size falling below the critical value of 100 individuals [77]. The maximum simulation time is an arbitrary period where persistence serves as a proxy for differential extinction rates under different parameters across time. That is, some populations will stably persist indefinitely (dash-dot line *Figure S2*), while others will eventually go extinct. However, the rate of their extinction may differ across settings as reflected by different rates of persistence across time. With a long simulation time, we ensure that the difference in persistence reflects the difference in the rate of extinction under different parameters (colored lines, *Figure S2*).

#### Controls

We generated two controls. (1) Drift, where the force driving allele frequency changes within populations is stochasticity due to finite population size, with *s*_max_ = 0. We define balanced polymorphism as the level of heterozygosity exceeding that under drift alone (*h*_model_/*h*_drift_ > 1.05). (2) Monomorphic populations, which evolve in the absence of genetic mutation. This control measures levels of survival under source-sink dynamics, i.e., under migration alone.

#### Implementation and Parameter Settings

For each parameter combination, we conducted 4,000 replicate simulation runs. We assume a constant carrying capacity of *K_i_* = 4,000 individuals in each population, a maximum simulation time of 100*K_i_*(400,000 generations), and a starting population size of 2,000 individuals in each population (V and R). We examine evolution under a wide range of magnitudes of varying selection, *s*_max_ = 0.0 (drift), 0.1, 0.15, 0.2, 0.25, 0.3, or 0.35 in combination with a period of the oscillating cycle, *P* = 12, 24, or 48, and migration rates *m,* where at carrying capacity the number of reciprocal migrants is equal to *M_i_* ∈[1,100], in increments of 1. For the simulations where the magnitude of fitness oscillations increases over time, *s*_max_ increases from 0.2 to 0.3 throughout the simulation, i.e. *β* = 0.1. The mean environmental effect in the harsh habitat was *S*_V_ = 0.3, 0.35, 0.4, or 0.45, with *S*_R_ = 0.0, and the proportion of fitness oscillations in R, *C* = 0, 0.167, 0.33, 0.5, 0.667, or 1. Recurrent mutation occurs at a rate of *μK_i_* = 0.05 per generation. In both populations *δ_i_* = 0.5 within the logistic growth model.

## Results

First, we provide an intuition on how genotypic diversity promotes the persistence of small populations in temporally varying environments, followed by the results from stochastic simulations.

Consider the change in the size of the subpopulations of carriers for each genotype, *d* and *a*, in V population under the eco-evolutionary population dynamics given in (1) and (2):

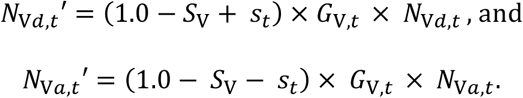

Recall, *s_t_*is a selection parameter, a genotype-based relative fitness deviation at time *t* due to varying selection. The ecological parameters are the environmental harshness *S*_V_ and density-dependent population growth rate *G*_V*,t*_. The expected change in population size due to the selection on carriers of *d* allele is then

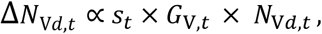

while due to the change in the number of *a* carriers

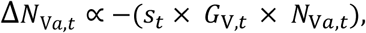

with *p*_V,*t*_ being the frequency of the ancestral allele, *a*, and *q*_V,*t*_ being the frequency of the derived allele, *d*, where *N*_V*d*,*t*_ = *N*_V,*t*_ × *q*_V,*t*_ and *N*_*ia*,*t*_= *N*_V,*t*_ × *p*_V,*t*_= *N*_V,*t*_ (1 - *q*_V,*t*_). Then, the change in population size, Δ*N*_V,*t*_, due to varying selection is a sum of changes in the number of individuals due to selection on *d* and due to selection on *a*:

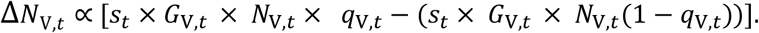

The smallest expected absolute change in population size due to selection, Δ*N*_V,*t*_ ∝ 0, occurs when *q*_V,*t*_= 1 - *q*_V,*t*_, i.e. *p*_V,*t*_ = *q*_V,*t*_= 0.5. On the other hand, the largest expected absolute change in population size due to selection, Δ*N*_V,*t*_ ∝ |*s*_*t*_ × *G*_V,*t*_ × *N*_V,*t*_| occurs when *q*_V,*t*_= 1 or 0, i.e. in a monomorphic population. The degree of variation in population size is, thus, a decreasing function of genetic variance (heterozygosity), 2*p*_V,*t*_*q*_V,*t*_.

For a small population at a size equilibrium, over a cycle of environmental variation, less perturbation in population size is expected to occur in a polymorphic population compared to a monomorphic population with the same parameters, which results in a smaller risk of reaching the absorbing state of extinction. The balanced polymorphism due to the spatial storage effect will therefore allow populations at a small equilibrium value (small immigration rate) to stably persist when they would perish in the absence of evolution. Here, rapid evolution occurs due to a rapid change in the sizes of subpopulations with differing genotypes.

While one subpopulation rapidly declines, the other counterbalances this effect by a rapid increase, thereby stabilizing the overall population size and enabling smaller populations to persist at a higher rate than would be possible in the absence of diversity. This phenomenon is what we refer to as ‘rapid rescue’.

For illustration, we present total population size, *N*_V_, genotype subpopulation sizes, *N*_V*a*_ and *N*_V*d*_ under our eco-evolutionary model, which allows stochastic size fluctuations, and where the storage effect was detected (*Figure 1*). While each genotype subpopulation oscillates notably and intermittently drops to near extinction, their mirrored oscillations balance each other, resulting in milder total population fluctuations. This stability maintains total population sizes, which oscillate around 1,000 individuals, demonstrating the mechanism behind the rapid rescue described above.

**Figure 1.**
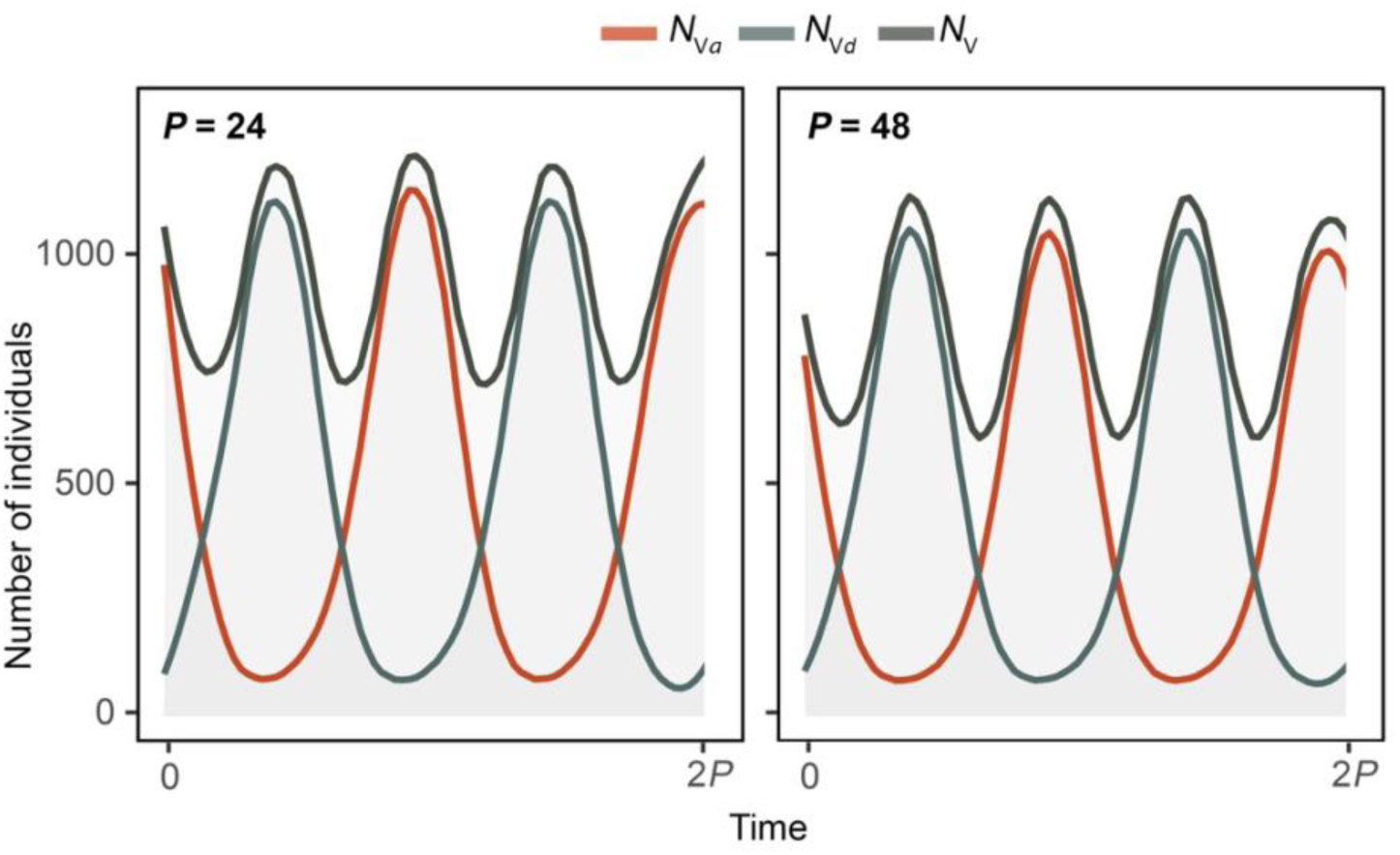
The expected number of individuals that carry the ancestral (*N*_V*a*_) or derived (*N*_V*d*_) allele over 2*P* generations in stably oscillating V populations of total size *N_V_*, where the storage effect was established (*h* = 3.40, and 3.90 for *P* = 24 and 48) under *P*×*s*_max_× *M*_V_ of 24 × 0.25 × 85 and 48 × 0.15 × 55. Simulations were conducted over 4,000 replicate runs with a starting size of 2,000 individuals and a carrying capacity (*K_i_*) of 4,000. The number of individuals (*N*_V_, *N*_V*a*_, *N*_V*d*_) was measured after 399,000 generations of density-regulated varying selection in 3,838 and 3,813 persisting populations for P = 24 and 48.

Under stochastic simulations, we find that the spatial storage effect arises in small populations of variable size experiencing temporal variation across a range of simulated parameters (*Figure S3*). In turn, the spatial storage effect increases the chance of persistence and average survival time while lowering the minimum number of migrants required to sustain a population compared to migration alone (source-sink dynamics) (*Figures 2* and *3, Figure S4*). The benefits of the storage effect are maintained when there is an increase in the magnitude of fitness oscillations over time (*Figure 4*) or through the introduction of temporally varying selection in the refuge, which reduces the level of spatial heterogeneity across the landscape (*Figure S5*). These results suggest that the spatial storage effect promotes rapid evolution and rapid rescue in populations inhabiting heterogenous temporally varying environments. We give details in the subsections below.

**Figure 2.**
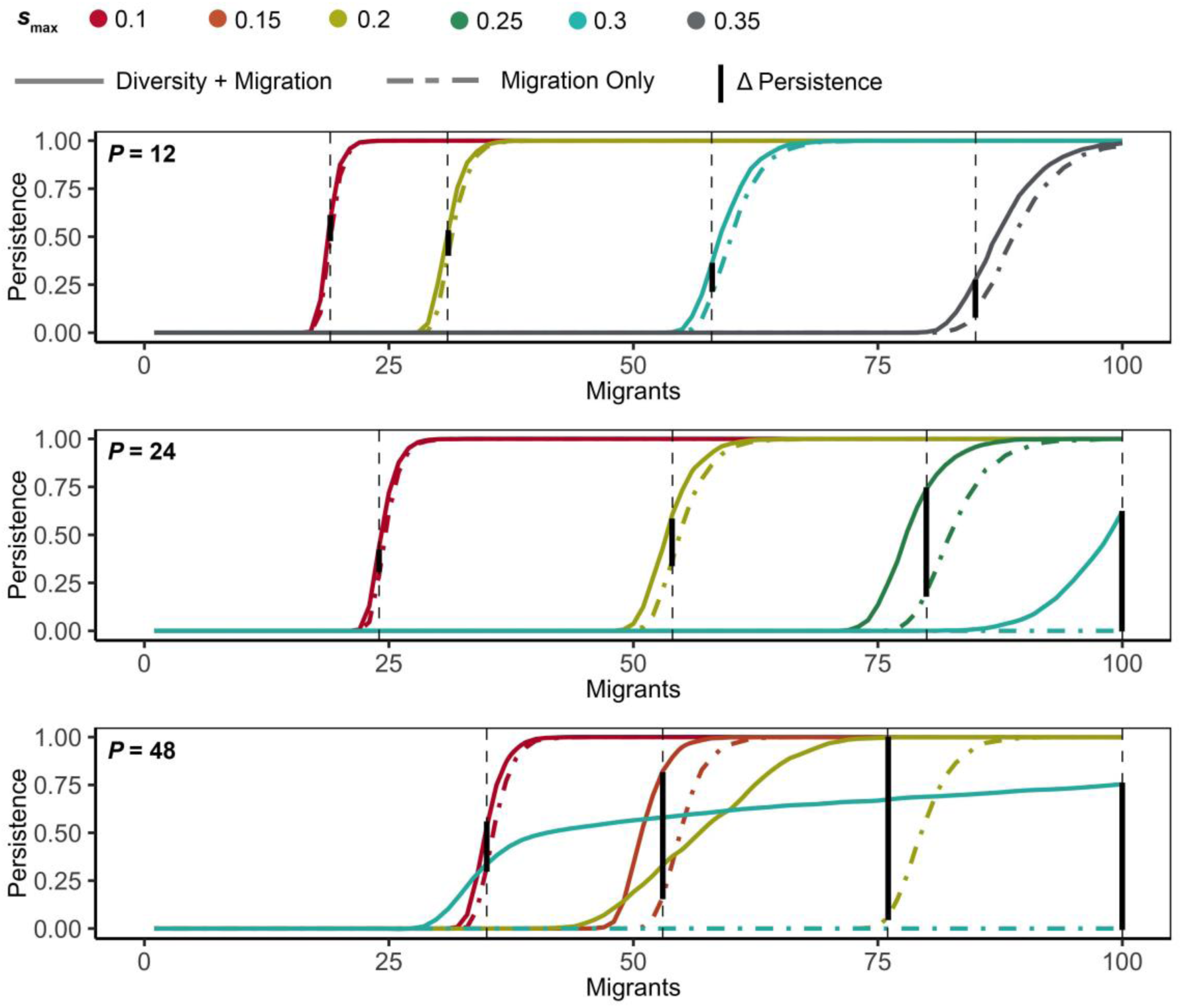
Population persistence for at least 100*K_I_* generations in a harsh habitat that is under temporally varying selection. Simulations assumed *P* = 12 (top), *P* = 24 (middle), and *P* = 48 (bottom), with *s*_max_ = 0.1 (red), 0.15 (orange), 0.2 (light green), 0.25 (green), 0.3 (turquoise), and 0.35 (grey). The environmental mean effect was *S*_V_ = 0.35. Dashed lines represent monomorphic scenarios (migration only), while the solid lines represent scenarios that allow recurrent mutation with *μ* = 0.0000125 (diversity and migration). Simulations were conducted over 4,000 replicate runs with a starting size of 2,000 individuals and a carrying capacity (*K_i_*) of 4,000. The solid black vertical lines showcase the increase in persistence between the model (migration + diversity) and control (migration only). These lines demonstrate that the storage effect led to the rapid rescue of these populations by reducing the minimum number of migrants required for the same level of persistence.

**Figure 3.**
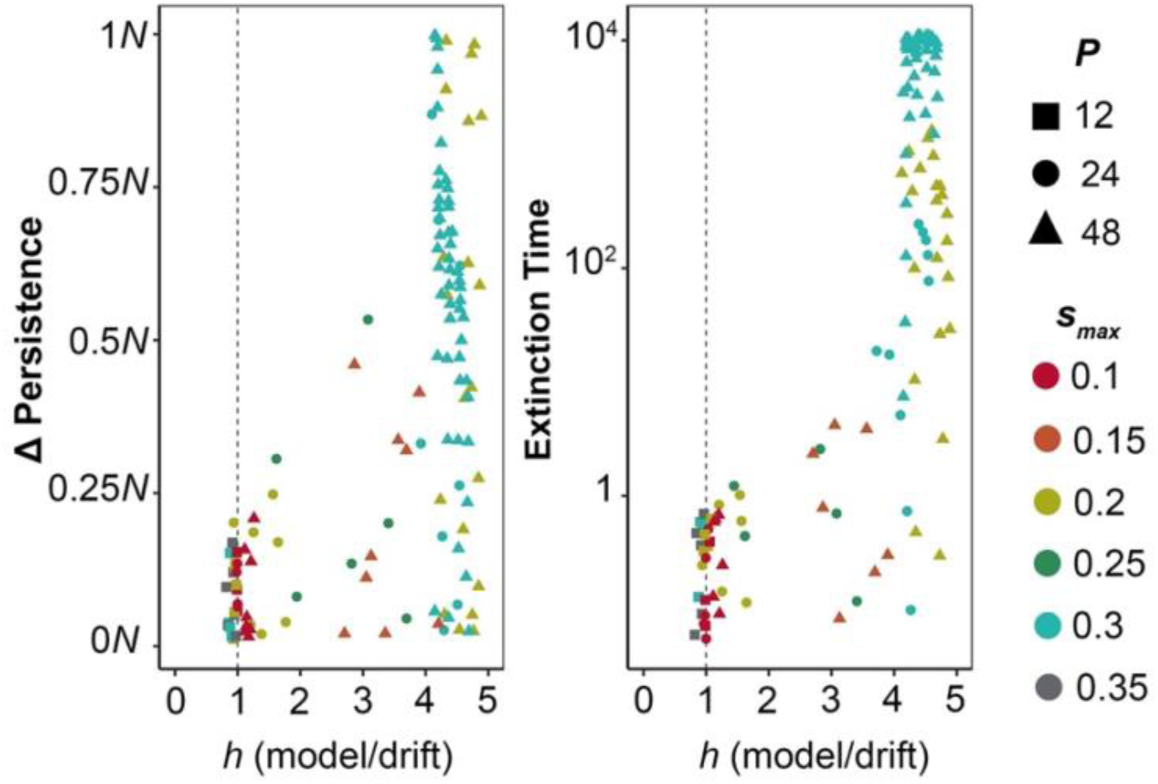
Impact of the level of genetic polymorphism on the differential persistence and the time at which a population became extinct (extinction time). Left panel: difference in population persistence (polymorphic populations compared to monomorphic controls). Right panel: expected extinction time for populations with differential persistence, measured as the relative increase in expected extinction time in the test populations compared to monomorphic controls (test-control)/control), note non-linear scale display on the y-axis. Simulations assumed a variety of *S* (*S* = 0.3, 0.35, 0.4, 0.45) and *s*_max_ (*s*_max_ = 0.1, 0.15, 0.2, 0.25, 0.3, and 0.35) with *P* = 12, *P* = 24, and *P* = 48, and were conducted over 4,000 replicate runs in temporally varying environments with a starting size of 2,000 individuals, carrying capacity (*K_i_*) of 4,000, and recurrent mutation rate *μ* = 0.0000125. We included test populations with at least 5% higher rate of persistence compared to the control populations to avoid a cluster of points around zero in cases where migration alone results in full persistence. A high level of persistence and a high average survival time appear to arise when larger levels of genetic polymorphism exist, with values to the right of the vertical dashed line falling under the spatial storage effect.

**Figure 4.**
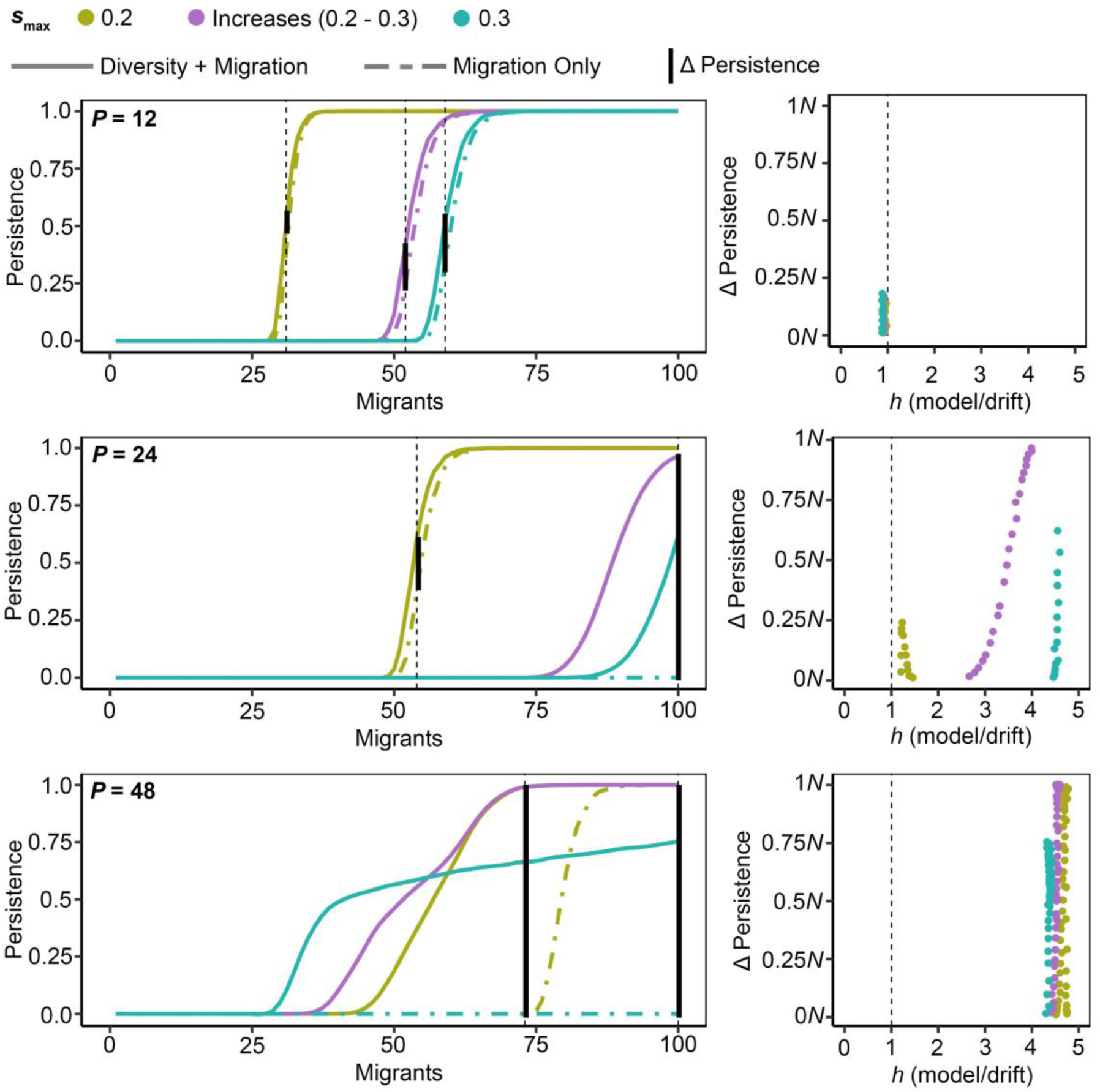
The population persistence in a habitat that is under temporally varying selection for at least 100*K_i_* (400,000) generations. Simulations assumed *P* = 12 (top), *P* = 24 (middle), *P* = 48 (bottom), each with *s*_max_ = 0.2 (light green), increasing from 0.2 to 0.3 (purple), or 0.3 (turquoise), with *S*_V_ = 0.35. The left panels show the persistence for test populations under the recurrent mutation with *μ* = 0.0000125 (solid line) or under the control (migration alone, dashed line). The solid black vertical lines showcase the increase in persistence between the two. The right panels show the level of persistence given the heterozygosity level. Simulations were conducted over 4,000 replicate runs with a starting size of 2,000 individuals and a carrying capacity (*K_i_*) of 4,000 in each subpopulation. Note that the populations that experienced a gradual change in selection (*s*_max_ = 0.2 -initial- to 0.3) had higher rates of persistence compared to the populations that were immediately placed under a high level of selection (*s*_max_ = 0.3).

### The spatial storage effect generates balanced polymorphism that leads to rapid rescue of small populations in temporally varying environments

Population genetics theory suggests that the spatial storage effect arises in finite constant-size populations in proportion to population size and the magnitude of varying selection, but inversely proportional to the rate of environmental change [51]. Here, we modeled rapidly changing conditions that emulate seasonal changes in environments, where a population experiences a season for six, twelve, or twenty-four discrete generations (*P* = 12, 24, or 48), but under stochastically varying population sizes. In addition, we focused on populations with a limited influx of migrants, where migration is not likely to ensure absolute persistence. We find that the spatial storage effect arises under reduced density dependence in small populations, and follows patterns reported by population genetics theory [51]. In particular, previous research suggests that in rapidly shifting environments, such as *P* = 12, the spatial storage effect would be more effective in populations approaching panmixia. Indeed, we find that under *P* = 12, balanced polymorphism is absent (*Figure S3*), likely due to limited migration. However, as *P* increases, so do the levels of balanced polymorphism, as proposed by population genetics findings [51], resulting in the spatial storage effect arising under a broad set of parameters (*Figure S3*), despite observed relatively small and stochastically changing population sizes.

The spatial storage effect increases population persistence across a wide range of scenarios. With rapid environmental change (*P* = 12), genetic diversity is maintained solely through the influx of mutants resulting in only a modest increase in persistence compared to the monomorphic control (*Figure 2*; examples shown by black vertical bars). However, as *P* increases and balanced polymorphism becomes more prominent, population persistence notably improves. This implies that the spatial storage effect results in an increase in population persistence across a range of parameters, including the rate of environmental change (number of generations that experience environmental oscillations, *P* = 24 or 48), the magnitude of varying selection (*s*_max_), the number of migrants exchanged between populations (*M*), and a range of harshness in habitats (*S*_V_) (*Figure 2, Figure S4*). The effect is larger with higher levels of balanced polymorphism (*Figure 3*, refer to *Figures 2* and *S3*). For example, with *P* = 24 when the number of migrants approaches a rate equivalent to 100 migrants, and with *P* = 48 with a migration rate equivalent to about 50 or more migrants, under carrying capacity, we observe notable increases in population persistence. Under these conditions, we observe that all populations (*Figure 2*, light green line) or a considerable proportion of populations (*Figure 2*, turquoise line) persist until the maximum simulation time for *P*=48, while all populations under source-sink dynamics (migration only) become extinct at the same settings (migration rate).

When comparing populations with genetic polymorphism (diversity and migration) to monomorphic populations (migration only), the spatial storage effect leads to an increase in population persistence at the same migration rates compared to the populations under source-sink dynamics alone. Over the range of parameters, therefore, we observe a notable reduction in the minimum number of migrants needed for a population to persist at the same rate in test populations compared to control populations (*Figure 2*, *Figure S4*). The conditions for persistence are thus broadened due to rapid rescue.

Additionally, there is a clear trend of increased average survival time before extinction with high levels of heterozygosity, serving as a proxy for the strength of the storage effect in a population (*Figure 3*). Even when populations did not reach the maximum simulation time, they still survived for longer than those under the source-sink dynamics alone. This is an important result of the storage effect since a longer survival time could provide more opportunities for the environmental conditions to change or for the population to expand their range into a habitat that is more favorable, thus ultimately avoiding extinction. Therefore, the spatial storage effect may also act to provide more time for populations to respond in other ways that may ultimately lead to population endurance.

### Robustness of rapid rescue to increasing magnitude of varying selection and decreasing levels of spatial heterogeneity

Next, we modeled the incremental increase in the magnitude of environmental oscillations (*s*_max_ = 0.2 → 0.3) over the course of simulations as driven by global climate change [78] (*Figure 4*). Here, the increase in the magnitude of varying selection produces patterns of persistence at intermediate levels compared to the starting and ending *s*_max_, for the test and control populations, as well as corresponding intermediate levels of heterozygosity (*Figure 4*, right panels). Hence, in the cases where the spatial storage effect leads to a sizable increase in persistence at constant *s*_max_, it also leads to a proportional increase in persistence under increasing *s*_max_. In other words, the level of persistence for the populations that experience a gradual change in their environment over time (*s*_max_ = 0.2 to 0.3) are higher than for those that experience a harsh shift immediately (*s*_max_ = 0.3). These results indicate that populations will be more likely to persist for a longer time when the magnitude of environmental oscillations increases gradually than during the sudden onset of strong environmental oscillations, such as migration to a new variable habitat or during an extreme climate event.

Lastly, we decrease the level of spatial heterogeneity by introducing temporally varying selection to the R population, with the magnitude of *Cs*_max_. Despite the reduction in spatial heterogeneity, an increase in population persistence is still evident, but to a lesser degree than when *C* = 0. The relative increase in persistence gradually diminishes as *C* increases from 0 (control) to 0.1667 and 0.33 (*Figure S5*). Notably, this pattern disappears when *C* ≥ 0.5, as both the test populations (diversity and migration) and the control populations (migration alone) fail to persist. In general, increasing *C* creates harsher conditions and the opportunities for population persistence rapidly shrink (the minimum number of required migrants increases) for both test and control settings. Overall, when a low level of environmental variation is introduced into the refuge, rapid rescue may occur but to a lesser degree than with high levels of spatial heterogeneity (*C* = 0.0).

In conclusion, the storage effect broadens the range of conditions under which populations of stochastically variable sizes can persist in the face of rapidly changing conditions by reducing the number of migrants required for persistence compared to source-sink dynamics alone.

## Discussion

In this study, we investigated whether the spatial storage effect can arise in finite populations inhabiting spatially heterogeneous variable habitats under relaxed density dependence and whether it can promote their persistence. Indeed, the spatial storage effect arises and promotes population persistence beyond levels expected in the absence of balanced polymorphism, thus leading to rapid evolutionary rescue in rapidly changing environments. The spatial storage effect can arise as both competing genotypes remain present in populations as environments change. The rapid population decline due to selection against detrimental genotypes is offset by a rapid increase due to selection favoring beneficial genotypes, across all environments, resulting in rapid rescue.

The theory of the storage effect has a long tradition in community ecology, with recent extensions into population genetics. The concept of the storage effect in the context of species coexistence was pioneered by Peter Chesson [56,79–81]. The theory proposes that environmental variability can promote coexistence in competitive systems under heterogenous varying selection, by enabling less abundant species to capitalize on favorable conditions while being partially buffered from detrimental conditions as environments change. Recently, the storage effect has been adapted to population genetics, emerging as a key mechanism that explains the existence of temporally varying balanced polymorphism. For example, heterogeneous varying selection may arise across life stages with reduced effect in overlapping generations [57,82], in the presence of phenotypic plasticity [52], under maternal effects [53], or when the magnitude of varying selection differs across spatial gradients (creating the so-called spatial storage effect) [51]. Gulisija and Kim (2015) found that the spatial storage effect promotes long-lived balanced polymorphism across a wide range of naturally plausible parameters in competitive systems modeled within the Wright-Fisher population framework [51]. More recently, Y. Kim (2023) extended this work to incorporate eco-evolutionary dynamics, proposing that correlated oscillations in both carrying capacity and allelic fitness may produce outcomes that differ from traditional population genetics predictions [32]. While this work was groundbreaking in incorporating the spatial storage effect into an eco-evolutionary framework, it did not examine stochastic perturbations to population size or explore the possibility of population extinction.

Our study incorporates demographic stochasticity, relaxing density dependence while allowing for extinction. This framework enables an evaluation of the role of the spatial storage effect in population persistence— an aspect that had not been previously examined, despite the recognition that balanced polymorphism is critical for adaptation to changing conditions [34]. We find that the storage effect not only promotes population persistence under eco-evolutionary dynamics but also exhibits the same key population genetics properties observed in constant-size Wright-Fisher models, under constant carrying capacity. In this scenario, expected oscillations in allele frequencies are captured by density-regulated selection. This suggests that since the balancing genetic effects of the spatial storage effect are preserved between the Wright-Fisher population and the eco-evolutionary dynamics examined here, the principle of rapid rescue may extend to other scenarios where other forms of the storage effect generate balanced polymorphisms, which were previously explored only in the framework of population genetics theory [49–56].

Various forms of the storage effect may offer naturally plausible mechanisms that facilitate evolutionary rescue in temporally varying environments. Two main processes that operate in populations are population size fluctuations and evolutionary change, yet they tend to be separated in theoretical studies [33]. Without examining the two processes jointly, we cannot consider how genetic adaptation rescues populations otherwise destined to perish, a phenomenon known as evolutionary rescue [60,62,77]. The theory of evolutionary rescue provides a framework that incorporates demographical and evolutionary dynamics in populations whose sizes vary [65,66,83–88]. Here, we demonstrate that evolutionary rescue can occur via the spatial storage effect while operating within small populations that occupy temporally varying habitats. In addition, we propose that a population exposed to a gradual increase in the magnitude of environmental variation has a higher chance of undergoing evolutionary rescue than the populations placed immediately into drastically changing conditions.

Despite evolutionary rescue’s relevance to populations facing rapidly changing environments, few studies have explored evolutionary rescue under temporally varying selection [e.g. 89]. However, this connection needs to be explored since conditions that change quickly may necessitate rapid adaptation from balanced genetic polymorphism. To predict the fate of future populations that are at risk for extinction due to increasingly extreme shifts driven by climate change [11,78,90,91], a synthesis of models of balancing selection and those of population size dynamics in rapidly changing environments is essential. Here, we propose a model that utilizes stochastically varying population sizes and incorporates the spatially heterogenous temporally varying selection to examine conditions for rapid rescue in the face of rapid environmental change.

A caveat of this study is that we only examine one form of the storge effect, the spatial storage effect. The spatial storage effect relies on strong spatial heterogeneity across an environment and reciprocal migration. As a proof-of-concept, we examined a scenario involving populations that exchange migrants proportionally to their sizes, which are allowed to stochastically vary. In this scenario, smaller populations in harsher environments may contribute disproportionately fewer migrants to refuge populations compared to migration under constant sizes. This likely limits the spatial storage effect, which requires notable migration, more so when environments oscillate rapidly [51]. How the other common forms of the storage effect, which do not rely on migration, impact eco-evolutionary dynamics in rapidly changing conditions is not clear. For example, the genomic storage effect arises under phenotypic plasticity [53] and does not depend on migration but on genetic recombination. Boarder forms of the storage effect need to be addressed in future studies in order to ascertain if the mechanisms offer a more widely applicable paths to rapid rescue.

In conclusion, our study expands our understanding of scenarios where the spatial storage effect can be established and uncovers a mechanism by which it promotes population persistence in the face of rapidly changing environmental conditions, a process we named rapid rescue. Rapid rescue could affect eco-evolutionary dynamics in other cases of environmental change. For example, populations may undergo range expansion or habitat invasions. Range limits are a major concern for populations under the environmental variation caused by climate change [20]. In addition, previous studies have demonstrated that past invaders and projected future climate migrants are particularly likely to come from habitats with temporally variability [23,92]. Studying evo-evolutionary dynamics in small founder populations originating from heterogeneous varying habitats could provide insights into how invaders establish in a novel habitat. Understanding the role of the storage effect in these other scenarios will become increasingly important as more populations face extreme environmental changes [11,78,90,91].

## Acknowledgments

We would like to thank the UNM Center for Advanced Research Computing, supported in part by the National Science Foundation, for providing the high-performance computing resources used in this work. We would also like to thank the members of the Gulisija Computational Genetics and Evolution Lab for their assistance throughout this study.

## Author Contributions

DG conceived the study and designed the initial model and simulation code. ENR modified the code and generated the data and figures. DG and ENR designed the study and co-wrote the article.

## Supplementary Material

**Figure S1.**
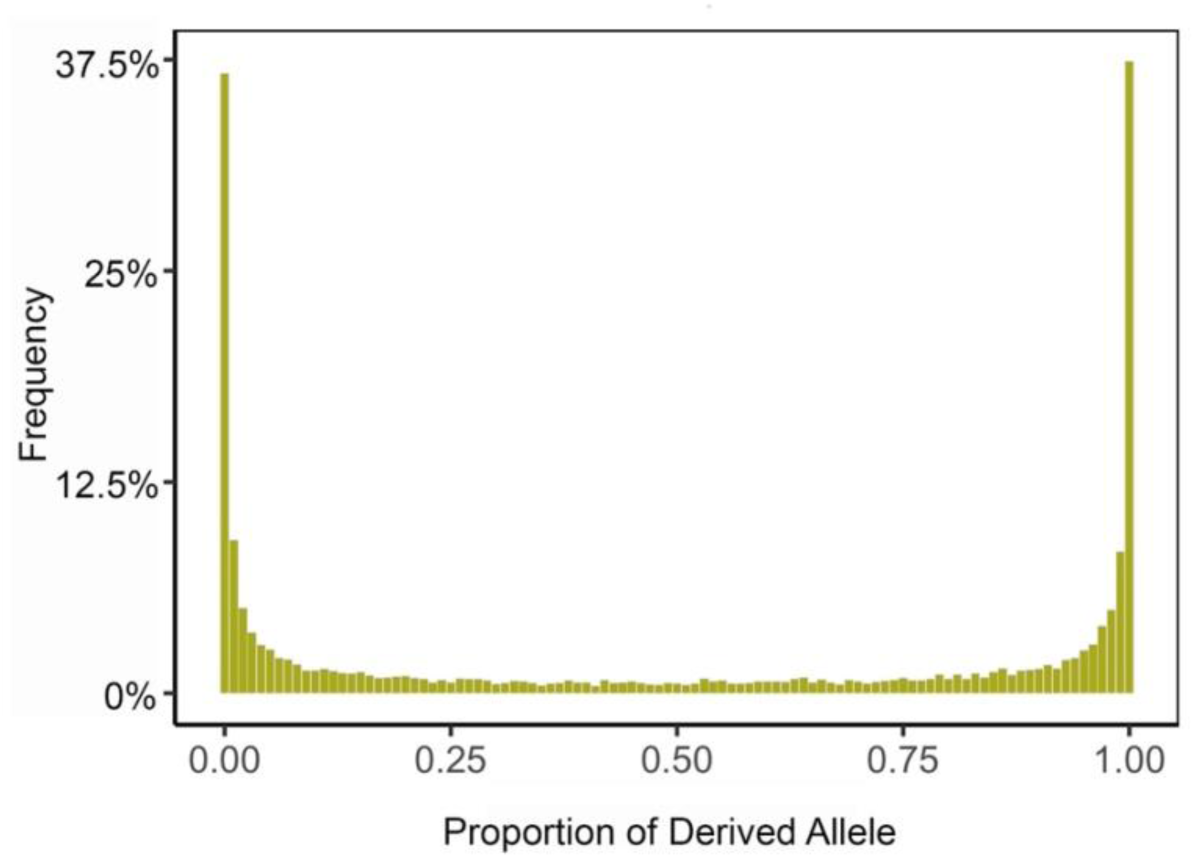
Distribution of the derived allele frequencies at the equilibrium between mutation and drift. The distribution was generated assuming a recurrent mutation rate *μ* = 0.0000125 and a starting population size at the carrying capacity of 8,000, and by conducting 16,000 replicate simulation runs with equilibrium confirmed at 1,280,000 generations. At the onset of varying selection, the starting allele frequencies in the two populations, V and R, were assigned by sampling from this distribution. For cases where the R population was at a lower magnitude of varying selection, compared to the V, distinct equilibria allele frequency distributions were generated. The generation of these distributions allowed all test simulations to start at drift-mutation (*C* = 0) or reduced selection-mutation (0 < *C* < 1) equilibrium.

**Figure S2.**
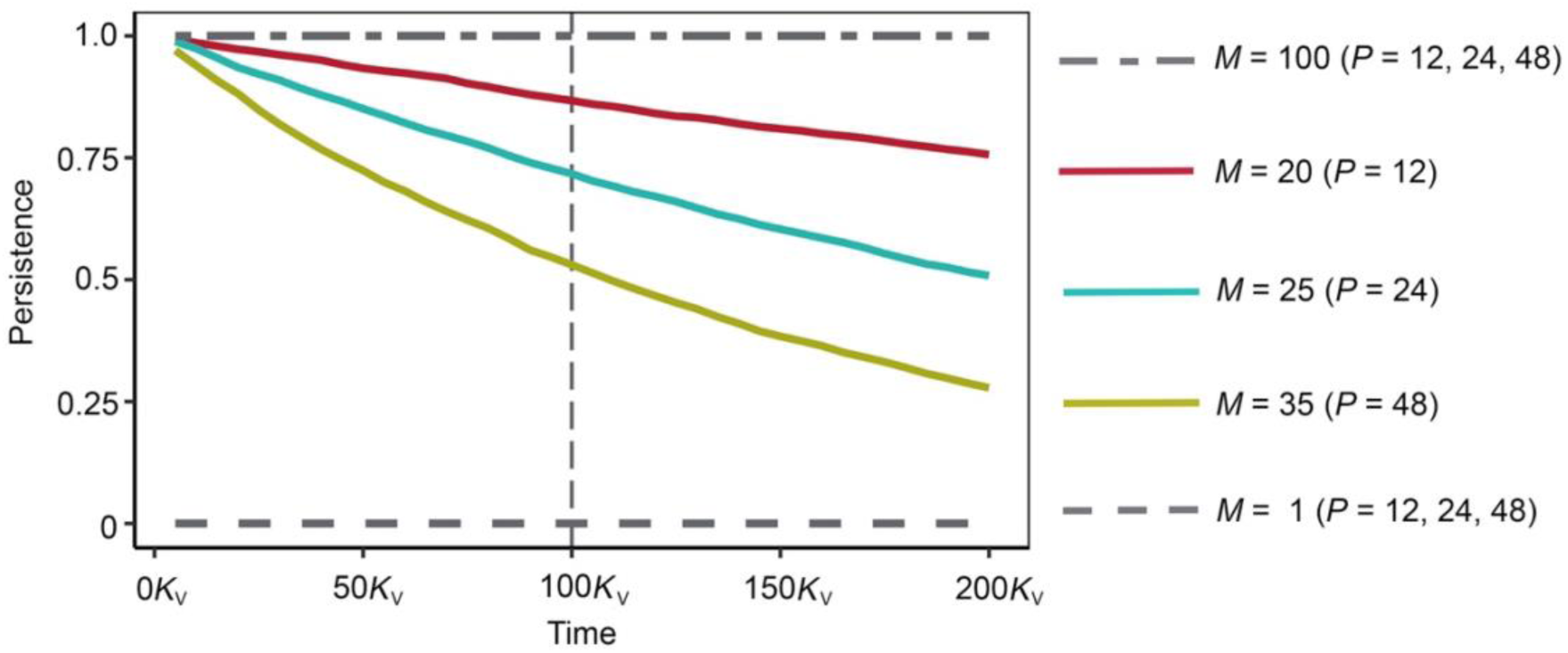
V population persistence over time under different eco-evolutionary scenarios. Simulation levels were tested across *P* values (red: *P* = 12, blue: *P* = 24, green: *P* = 48), and migration levels (low *M* = 1, intermediate *M*: 20/25/35 for *P* = 12/24/48 respectively, and high *M* = 100) with *s*_max_ = 0.1. The dashed line represents early extinction under low migration rates, and dash-dot lines represent stable persistence under high migration observed under all *P*s. Simulations were conducted with 4,000 replicates and initial population sizes of 2,000 and a carrying capacity of *K_i_* = 4,000 individuals in each (*i* = R or V). For populations that will eventually become extinct, persistence at 400,000 generations (100*K*_V_) serves as a proxy of the rate of persistence over time.

**Figure S3.**
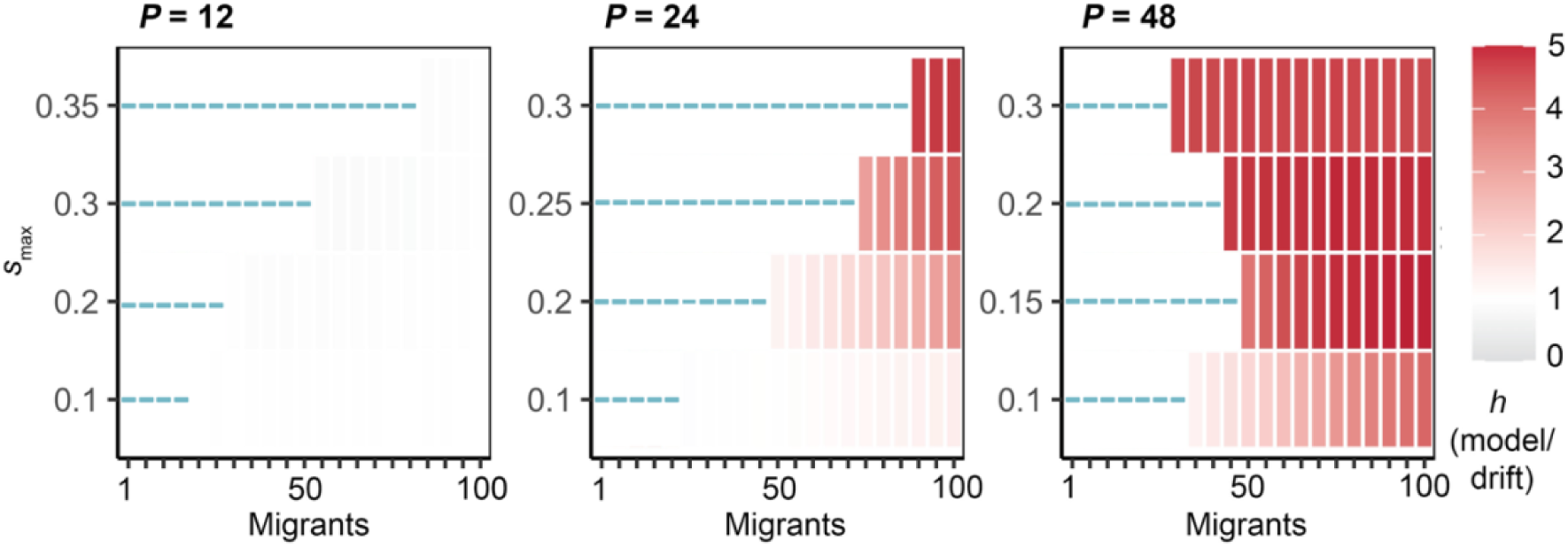
The expected levels of heterozygosity under eco-evolutionary settings where at least 5 percent of simulation runs reached persistence at 100*K*_i_ generations, under parameters given in Figure 1. The V populations where this threshold was not met (<5% persistence) are marked with a blue line. Simulations assumed *P* = 12, 24, 48 (left to right columns) and within each box, *s*_max_ increases from bottom to top, and the number of migrants increases from left to right, for all parameter combinations, with *S*_V_ = 0.35. We assume a constant carrying capacity of *K_i_* = 4,000 individuals in each population (*i* = R or V) with a starting size of 2,000 individuals. The spatial storage effect can be established in populations under a wide variety of conditions, including varying periods of temporal variation, number of migrants, and varying selection pressures. Moreover, the high levels of heterozygosity, indicating the establishment of the storage effect, correspond to parameter combination with a greater increase in persistence compared to the monomorphic control.

**Figure S4.**
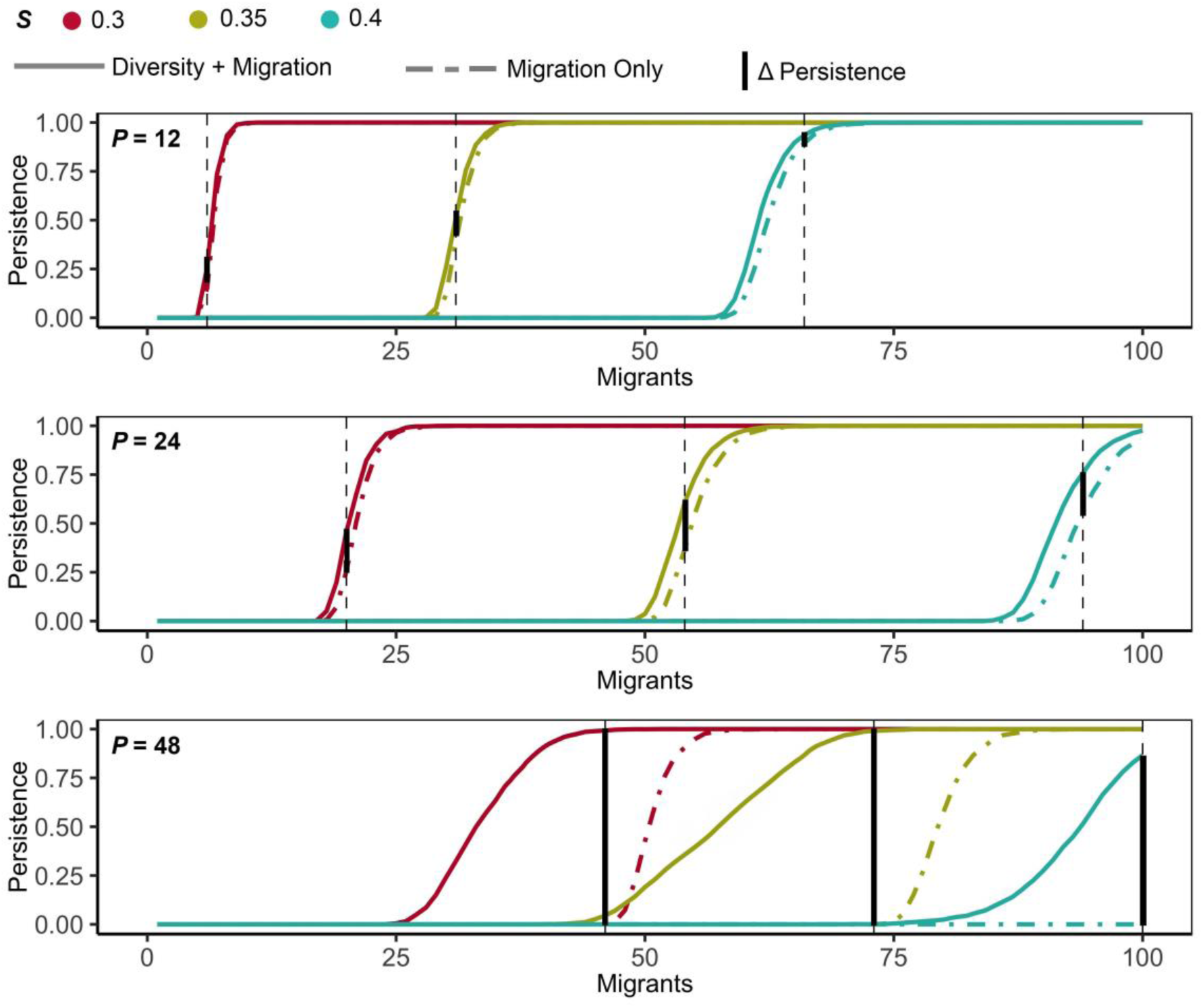
The proportion of V population that persisted for at least 400,000 generations under temporally varying environments across different degrees of environmental harshness (*S*_V_). Simulations assumed *S*_V_ = 0.3 (red), 0.35 (light green), 0.4 (turquoise) and *P* = 12 (top), *P* = 24 (middle), and *P* = 48 (bottom), with *s*_max_ = 0.2 for all parameter combinations. Simulations were conducted over 4,000 replicate runs with a starting size of 2,000 individuals and a carrying capacity of *K_i_*= 4,000 in both populations that exchanged migrants. Dashed lines represent monomorphic controls (migration only), while the solid lines represent simulations with recurrent mutation, *μ* = 0.0000125 (diversity and migration). The solid black vertical lines showcase the increase in persistence between the two, demonstrating that the storage effect rescued V populations across various environmental conditions.

**Figure S5.**
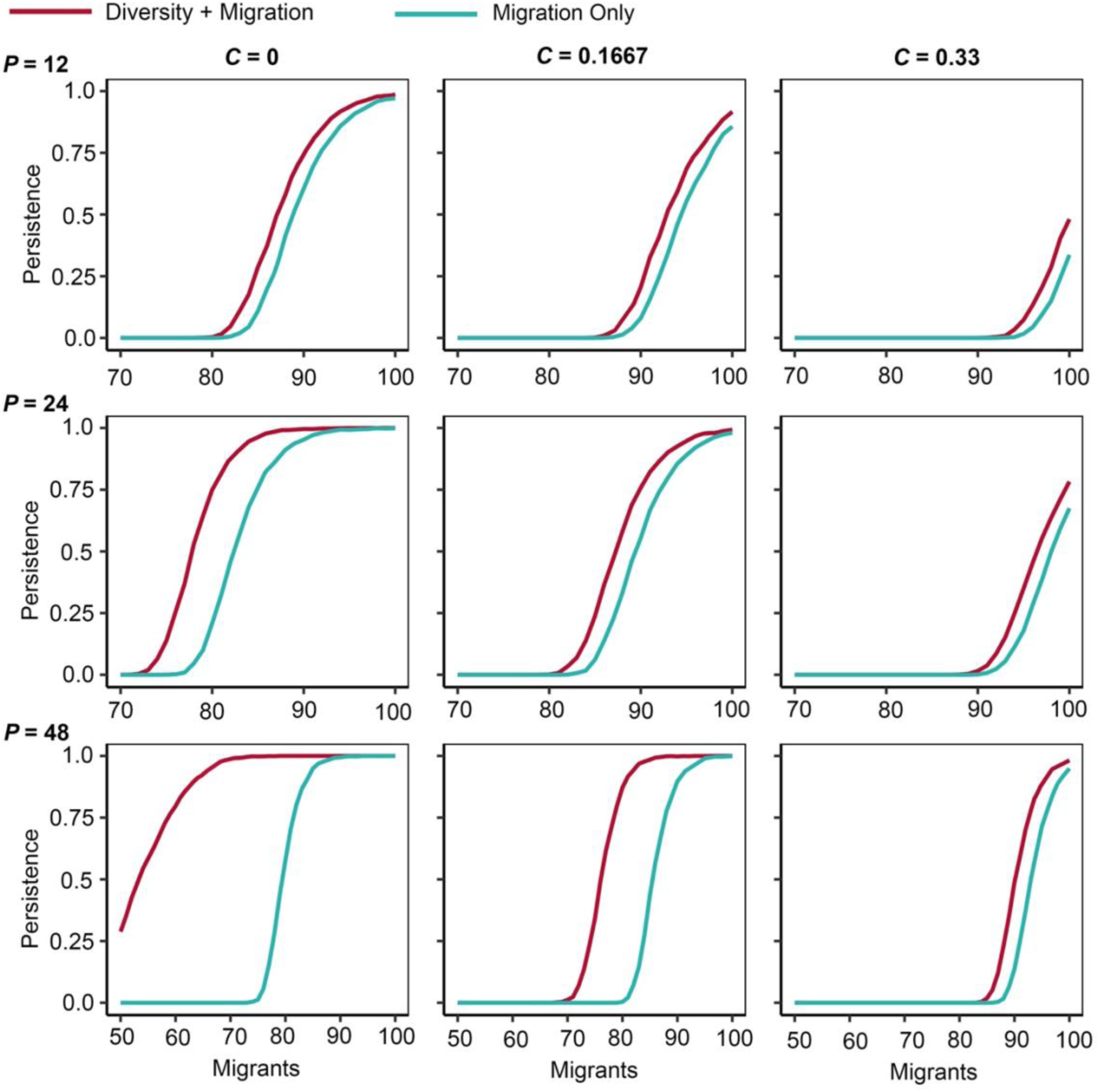
Populations persistence for at least 100*K_i_* (400,000) generations in V when both habitats experience varying selection. The left panels show populations with proportional variation *C* = 0, the middle shows *C* = 0.1667, and the right shows *C* = 0.33. The rows of panels correspond to *P* values (top: *P* = 12, middle: *P* = 24, and bottom: *P* = 48). Within each plot, the persistence rate of test populations, with *μ* = 0.0000125 (diversity and migration) is shown in red and the persistence rate of monomorphic populations (migration only) is shown in turquoise. Simulations were conducted over 4,000 replicate runs with a starting size of 2,000 individuals and a carrying capacity (*K_i_*) of 4,000 in each subpopulation. When the level of spatial heterogeneity across the landscape is reduced by introducing environmental variation into the refuge (R), the storage effect is still able to produce a higher rate of persistence compared to the control populations, although to a lesser degree than when *C* = 0.

